# Targeting the Alk4 pathway protects against age-related bone loss

**DOI:** 10.1101/2025.10.24.684408

**Authors:** David E. Maridas, Shek M. Chim, Kathryn M. Burton, Emily Van Doren, Laura Gamer, Kyle Kirshen, Eren Keles, Karla Avalos, Radhika S. Kethani, Vicki Rosen

## Abstract

Osteoporosis is a chronic age-related condition in which imbalanced activities between bone-forming osteoblasts and bone-resorbing osteoclasts lead to the progressive loss of bone volume and quality. While drugs that target osteoclastic activity have been developed, there remains a lack of efficient therapies that increase osteoblast number and function in aging bones. Here, we investigated if Activin, known to increase in the circulation with age, plays a primary role in bone loss associated with aging. We showed that, in mouse femurs, levels of Activin signaling progressively increased with age and strongly correlated with the loss of trabecular bone. Furthermore, mice lacking the type I receptor for Activin, namely Alk4, in osteoblast progenitors (Alk4 cKO mice) had increased trabecular bone acquisition, osteoblast number, and bone formation rate. In addition, Alk4 cKO male mice were protected against early age-related trabecular bone loss observed at 1 year of age. These results indicate that Activin signaling inhibits bone formation and osteoblast activity and is likely associated with osteoporosis. To further test this, we injected 2-year-old male mice with a ligand trap (Alk4-Fc) to capture circulating Activin. Alk4-Fc protected against loss of trabecular bone in femurs and L5 vertebrae. Interestingly, Alk4-Fc also prevented a decrease in muscle mass in gastrocnemius, quadriceps, and triceps suggesting that circulating Activins also play a role in sarcopenia. In summary, our preclinical mouse models reveal that circulating Activins play a primary role in age-related bone loss and can be efficiently targeted to alleviate osteoporosis and sarcopenia in aging mice.

**One Sentence Summary:** In this study, we discovered a new way of preserving trabecular bone mass in aging mice by inhibiting activity of Alk4 pathway.

## INTRODUCTION

Osteoporosis is a chronic disease characterized by decreases in bone volume and strength that lead to increased skeletal fragility. The progressive age-related bone loss that ultimately causes osteoporosis has been extensively studied in humans and animal models, and virtually every adult will experience slower bone formation with age^1–6^. This decrease in bone formation can be accelerated by sex steroid decline, chronic diseases and their treatments, inflammation, disrupted metabolism, persistent skeletal unloading, and substance abuse^7–10^. Yet, in all these instances, the biological mechanisms that initiate the decline in bone formation remain elusive. Discovering new biological targets associated with the natural decline in bone formation has the potential to identify novel therapeutic strategies to fight osteoporosis.

In the adult skeleton, bone formation is maintained by the activity of osteoblasts, cells specialized in the production and mineralization of the bone matrix. Ligands of the TGFβ superfamily including, TGFβs, BMPs, GDFs, and Activins, exert profound regulatory effects on the differentiation and function of osteoblasts via two pathways: the Smad1/5/8 pathway (activated by BMPs and certain GDFs) and the Smad2/3 pathway (activated by TGFβs, Activins, and GDFs). A large body of literature shows that activation of the Smad1/5/8 pathway increases the differentiation and activity of osteoblasts^11–19^. In contrast, the Smad2/3 pathway inhibits osteoblast differentiation and mineralization^20–24^. While these effects on osteoblasts have been well documented, the role of the Smad1/5/8 and Smad2/3 pathways in the aging skeleton, specifically in age-related bone loss, has been understudied.

In this study, we sought to determine if activation of these pathways change when osteoblast activity diminishes and bone is lost in maturing mice. Our data reveal a significant increase in Smad2/3 activation in the maturing skeleton likely caused by a rise in circulating Activins. Interestingly, deleting Alk4, the type I receptor for Activins in bone, is sufficient to reduce Smad2/3 activation, increase osteoblast number and bone formation, and slow age-related bone loss in adult mice. Finally, we show that treating 24-month-old mice with a pharmaceutical ligand trap for circulating Activins blocks further loss of bone mineral density (BMD) and bone volume while also protecting against muscle wasting. Collectively, our results suggest that targeting age-related increase in circulating Activins is a promising avenue for preventing bone loss and the progression of osteoporosis.

## RESULTS

### Smad2/3 activation is increased in aging bones

Initially, we wanted to determine when bone mass started to naturally decline with age. We performed microCT analysis on the distal femurs of 3-, 6-, 12-, and 24-month-old female C57BL6J mice, times points when changes in trabecular bone were previously reported^25, 26^. We found a steady decline in trabecular bone volume throughout the study with a significant decrease from 3 to 6 months of age (**Fig. 1A**). Next, we extracted proteins from the femoral trabecular space and performed western immunoblots for phosphorylated Smad1/5/8 (pSmad1/5/8) and Smad2 (pSmad2) to measure activation levels of the BMP and Activin/GDF/TGFβ pathways, respectively. Levels of pSmad1/5/8 started to decrease at 12 months of age while an increase in pSmad2 was detected as early as 6 months of age (**Fig. 1B**). Statistical analyses revealed that trabecular bone volume was directly correlated with levels of pSmad1/5/8 and inversely correlated with pSmad2 levels (**Fig. 1C**). Next, we asked whether the age-related changes in the levels pSmad1/5/8 and pSmad2 were influenced by levels of receptor or ligand expression. We performed qPCR on RNA samples extracted from the trabecular space microenvironment to assess the expression of receptors and ligands known to mediate Smad1/5/8 and Smad2 phosphorylation. The RNA expression of most ligands and receptors did not change with age (**Fig. 1D**). *Alk5* was the only receptor showing a significant decrease in expression at 12 and 24 months of age (**Fig. 1D**). Our results indicated that the age-related differences in pSmad1/5/8 and pSmad2 levels in bone were not caused by local change of receptor or ligand expression.

**Figure 1.**
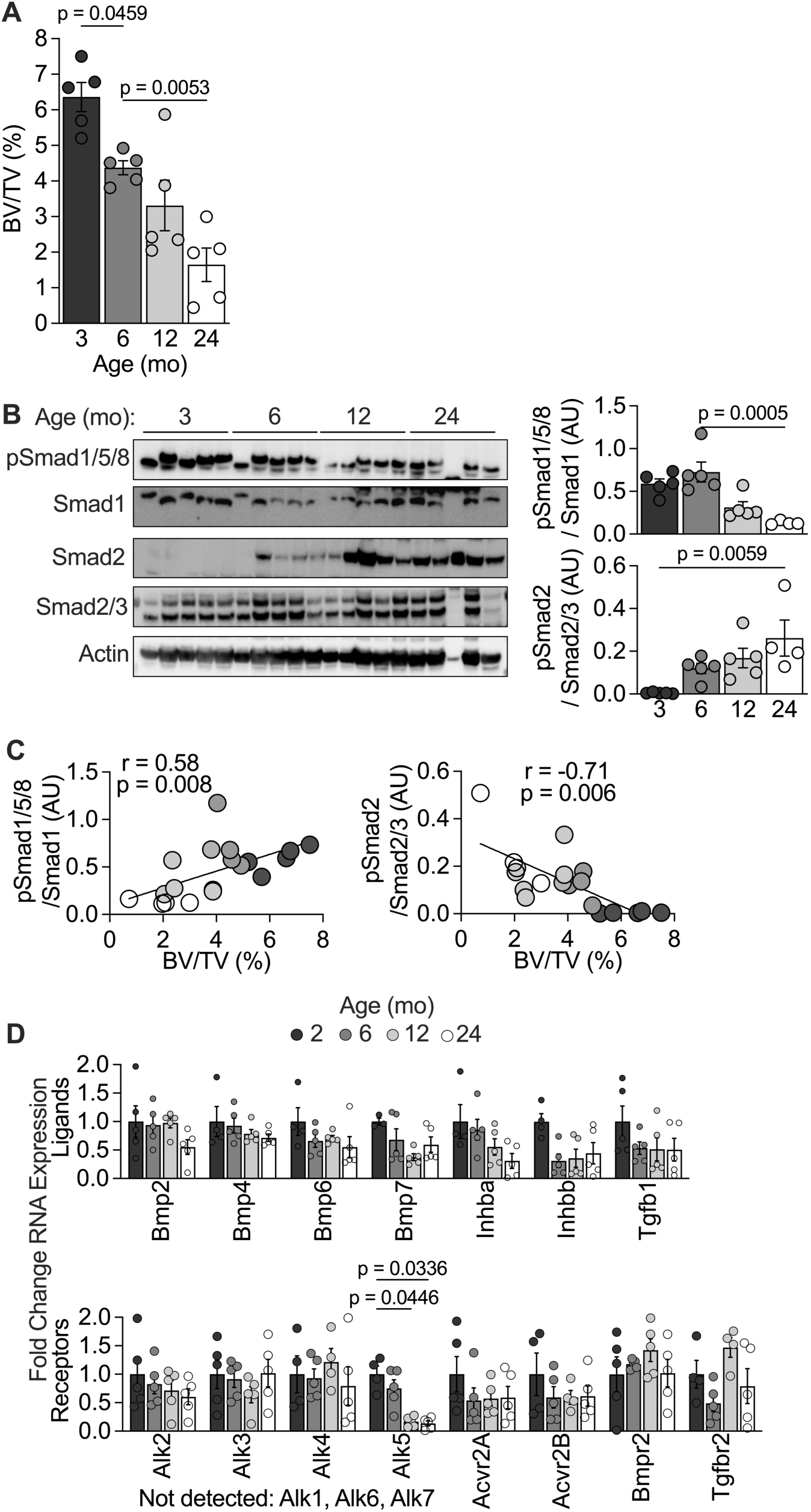
Smad1/5/8 and Smad2 activation levels in the bones of aging mice. (**A**) MicroCT measurements of trabecular bone volume / total volume (BV/TV) in the distal region of the femur of aging female C57Bl/6J mice. (**B**) Western Immunoblots and densitometry analysis of phosphorylated Smad1/5/8 (pSmad1/5/8), Smad1, phosphorylated Smad2 (pSmad2), Smad2/3, and actin performed on protein lysates extracted from the marrow of aging female C57Bl/6J mice. (**C**) Correlation coefficients and linear regression analyses between pSmad levels and BV/TV. (**D**) RNA expression of ligands and receptors of the TGFβ superfamily involved in phosphorylating Smad1/5/8 and Smad2 in femurs of aging mice. Statistical analyses include 1-way ANOVA (A, B), linear regression and correlation analyses (C), and 2-way ANOVA (D) with multiple comparisons, p-values shown when < 0.05

### Circulating ligands are responsible for the age-related increase in Smad2/3 signaling

We hypothesized that alterations in circulating ligands, rather than local changes, were responsible for the increased pSmad2 levels observed after 3 months of age. To test this, we treated bone marrow stromal cells (BMSCs) with sera from 3-, 6-, and 12-month-old mice and measured pSmad2 levels. Treatment with sera from 6- and 12-month-old mice caused a marked increase in pSmad2 level in BMSCs when compared to sera from 3-month-old mice (**Fig. 2A**). To test whether circulating TGFβ was responsible for the increase in pSmad2, we depleted TGFβ from the sera samples using an anti-TGFβ antibody (TGFβ Ab). Depletion of TGFβ resulted in a considerable reduction of pSmad2 in BMSCs (**Fig. 2B**). However, the age-related increase in pSmad2 levels observed in sera from aging mice was not affected by treatment with TGFβ Ab (**Fig. 2B**). Next, we measured levels of circulating ligands known to activate the Smad2/3 pathway, i.e., Activin A and TGFβ1, in the sera of 2-, 6- and 12-month-old mice. We found increased levels of Activin A in sera of older mice (**Fig. 2C**) and no change in TGFβ1 level (**Fig. 2D**). Taken together, our results suggest that circulating ligands other than TGFβ are likely responsible for the increase in pSmad2 levels in the bones of aging mice.

**Figure 2.**
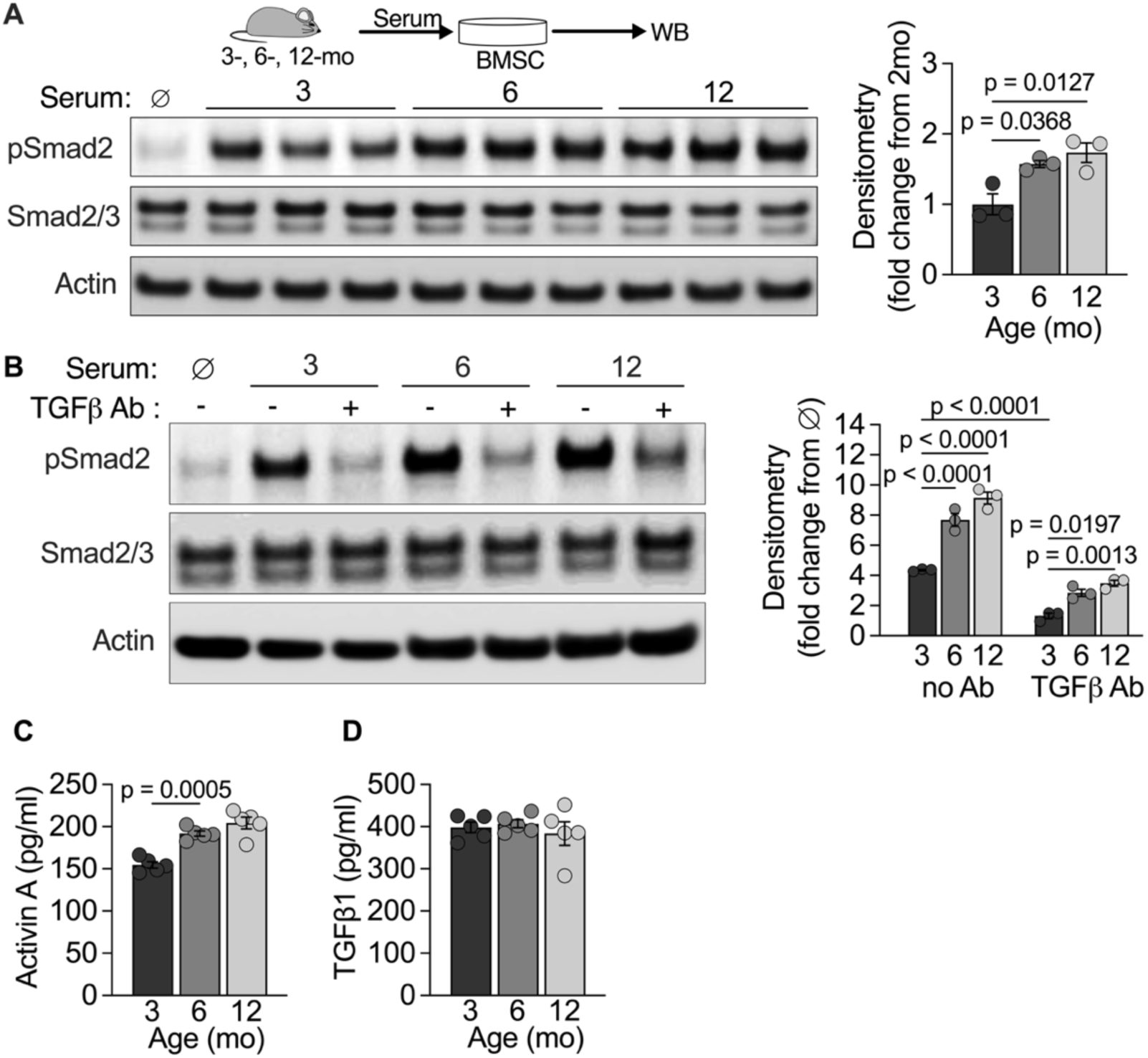
Smad2/3-activating ligands in the circulation of aging mice. Primary bone marrow stromal cells (BMSCs) cultures were treated with sera samples extracted from 3-, 6-, and 12-month-old female C57BL/6J mice. Proteins from BMSCs were extracted to perform western immunoblotting and densitometry analysis (**A**) of pSmad2, Smad2/3, and Actin. (**B**) The same experiment was performed with sera treated with anti-TGFβ antibody (Ab). Circulating levels of Activin A (**C**) and TGFβ1 (**D**) measured by ELISA in sera samples from 3-, 6-, and 12-month-old female C57BL/6J mice. Statistical analyses include 1-way ANOVA (A,C) and 2-way ANOVA (B) with multiple comparisons, p-values shown when < 0.05.

### Female Alk4 cKO mice have higher trabecular bone mass

Next, we hypothesized that blocking Smad2/3 activation by circulating ligands would affect bone mass and age-related bone loss in mice. In bone, circulating ligands bind Alk4 or Alk5 to phosphorylate Smad2/3. We previously showed that the expression of *Alk5* decreases considerably with age (**Fig. 1D**). This suggests that Alk4, the type I receptor for activins, is likely responsible for the increase in Smad2/3 activation levels in the mature skeleton. Thus, we deleted *Alk4* using Prx1-Cre (Alk4 cKO mice) to target osteoprogenitor cells that participate in the formation of trabecular bone. Despite *Alk4* being expressed in the developing limb bud^27^, Alk4 cKO mice were born with normal appendicular skeletons (**Supplemental Fig. 1A-B**) suggesting that signaling through this receptor is dispensable for embryonic skeletogenesis. Despite a significant decrease in *Alk4* RNA expression in the femurs of 4-week-old Alk4 cKO mice (**Supplemental Fig. 1C**), we did not detect any changes in femoral length (**Supplemental Fig. 1D**), trabecular bone volume (**Supplemental Fig. 1E**), or cortical bone area (**Supplemental Fig. 1F**) at this time point. In addition, femoral levels of pSmad1 and pSmad2 of 4-week-old Alk4 cKO mice were comparable to that of controls (Alk4^fl/fl^) (**Supplemental Fig. 1G**). Together, our results indicate that Alk4 expression in Prx1^+^ cells is not required for skeletogenesis and Smad2/3 signaling is likely provided by another type I receptor during skeletal development and early postnatal life.

As we observed increased levels of pSmad2 in bone after 3 months of age (**Fig. 1B**), we measured bone parameters of Alk4^fl/fl^ and Alk4 cKO mice by microCT at these time points. At 2 and 6 months of age, female Alk4 cKO mice had higher trabecular bone volume (**Fig. 3A**) and thickness (**Fig. 3D**) when compared to Alk4^fl/fl^. After 2 months of age, both Alk4^fl/fl^ and Alk4 cKO female mice exhibited significant loss of trabecular bone and by 12 months of age, there were no differences in trabecular bone parameters between genotypes (**Fig. 3A-F**). At the mid-diaphysis level, both cortical area and cortical thickness increased between 2 and 6 months of age and were not different in Alk4^fl/fl^ and Alk4 cKO female mice (**Fig. 3G-I**). Our data show that *Alk4* deletion in skeletal progenitors increases the amount of trabecular bone in young adult female mice.

**Figure 3.**
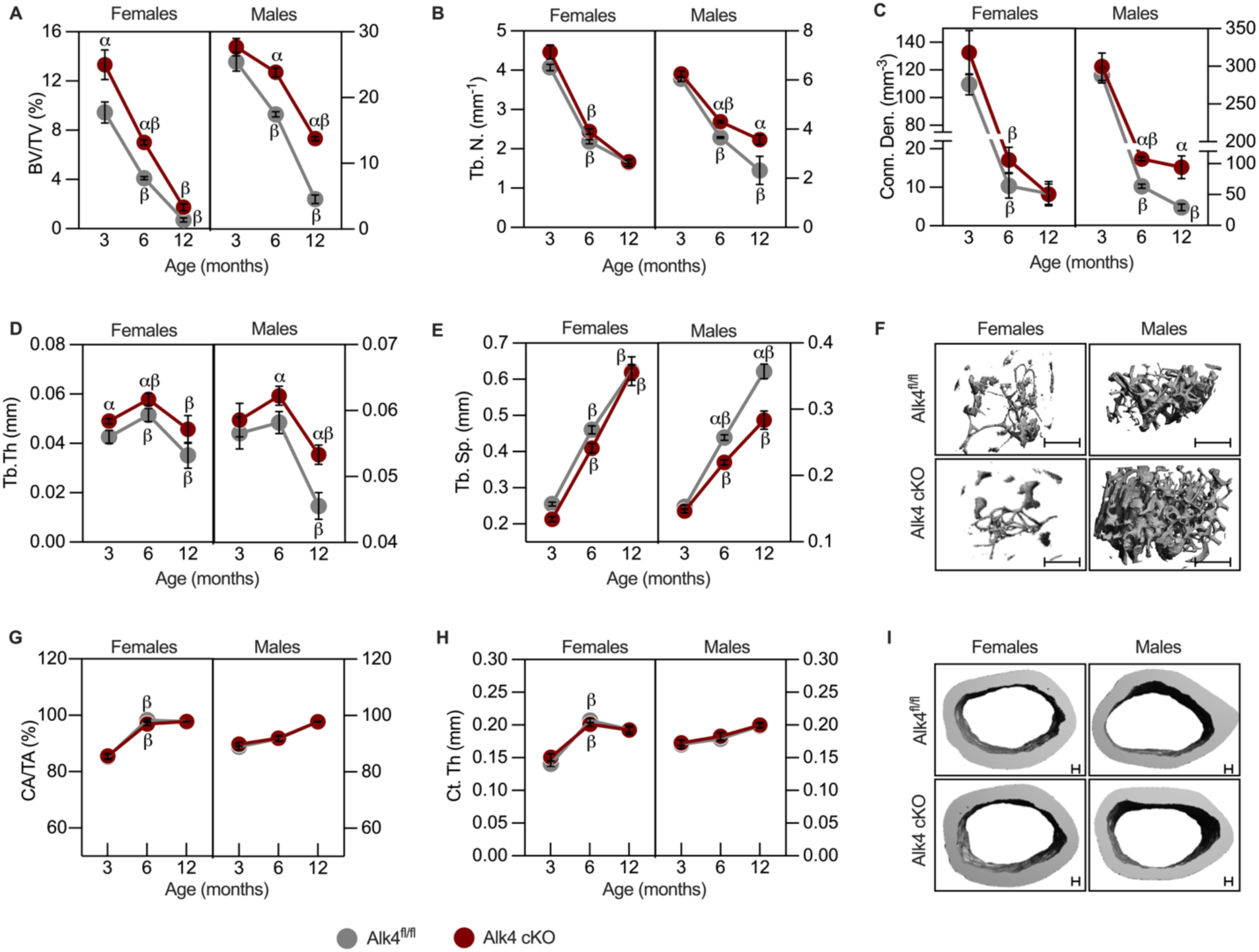
Adult Alk4 cKO mice have increased bone mass and are protected against early age-related bone loss. MicroCT parameters were measured in femurs of 3-, 6-, and 12-month-old Alk4^fl/fl^ and Alk4 cKO mice. Trabecular bone measurements were performed in the femoral distal region: (**A**) Bone volume / total volume (BV/TV), (**B**) Trabeculae number (Tb. N.), (**C**) Connectivity density (Conn. Den.), (**D**) Trabeculae thickness (Tb. Th.), (**E**) Trabeculae separation (Tb. Sp.), (**F**) Representative microCT images of trabecular bone at 12 months of age. Cortical bone measurements were performed at the femoral midshaft diaphysis: (**G**) Cortical area / total area (CA/TA), (**H**) Cortical thickness (Ct. Th.). (**I**) Representative microCT images of cortical bone at 12 months of age. n= 6-7 animals/genotype/sex/age group. Statistical analyses include 2-way ANOVA with multiple comparisons. α show adjusted p-value <0.05 when comparing between genotypes at the same time point. β show adjusted p-value <0.05 when comparing from a time point to the previous one within the same genotype.

### Male Alk4 cKO mice are protected against early age-related trabecular bone loss

At 2 months of age, there were no statistical differences between male Alk4^fl/fl^ and Alk4 cKO mice in all microCT parameters (**Fig**. **3A-E**). However, the rate of bone loss was markedly slower in Alk4 cKO males, and by 6 and 12 months of age Alk4 cKO males had higher trabecular bone volume (**Fig. 3A**), number (**Fig. 3B**), connectivity density (**Fig. 3C**), and thickness (**Fig. 3D**), and lower trabecular separation (**Fig. 3E**) when compared to Alk4^fl/fl^. At the cortical level, no differences were detected between Alk4^fl/fl^ and Alk4 cKO mice **(Fig. 3G-I)**. Together, our results indicate that age-related trabecular bone loss is slowed in Alk4 cKO male mice.

### Alk4 cKO mice have reduced Smad2 phosphorylation levels and increased osteoblastic activity

Since trabecular bone parameters were changed as early as 2 months of age in female Alk4 cKO mice, we investigated the levels of pSmad1/5/8 and pSmad2 in these mice. There was a significant reduction of pSmad2 levels in the trabecular space in femurs of Alk4 cKO female mice when compared to Alk4^fl/fl^ (**Fig. 4A**). We did not detect any change in pSmad1/5/8 levels between Alk4^fl/fl^ and Alk4 cKO mice (**Fig. 4A**). Next, we performed histomorphometric analyses on tibias to examine potential changes in bone cell number and activity in Alk4 cKO mice. There were marked increases in osteoblast number (**Fig. 4B**) and coverage (**Fig. 4C**) in the tibias of Alk4 cKO female mice compared to Alk4^fl/fl^. There was no difference in erosion surface (**Fig. 4D**), osteoclast coverage (**Fig. 4E**), or osteocyte number (**Fig. 4F**). Additionally, dynamic histomorphometry showed robust increases in both mineral apposition rate (**Fig. 4G**) and bone formation rate (**Fig. 4H**) in female Alk4 cKO mice. When we examined the levels of serum markers associated with bone resorption and formation, we found no difference in circulating levels of carboxy-terminal collagen crosslinks (CTX), a bone resorption marker, (**Fig. 4I**), but serum levels of N-terminal propeptide of type 1 collagen (P1NP), a marker for bone formation, were elevated in Alk4 cKO female mice compared to controls (**Fig. 4J**). Collectively, our data suggest that the increased trabecular bone mass observed in adult female Alk4 cKO mice results from a reduction in pSmad2 activity and higher osteoblastic number and activity.

**Figure 4.**
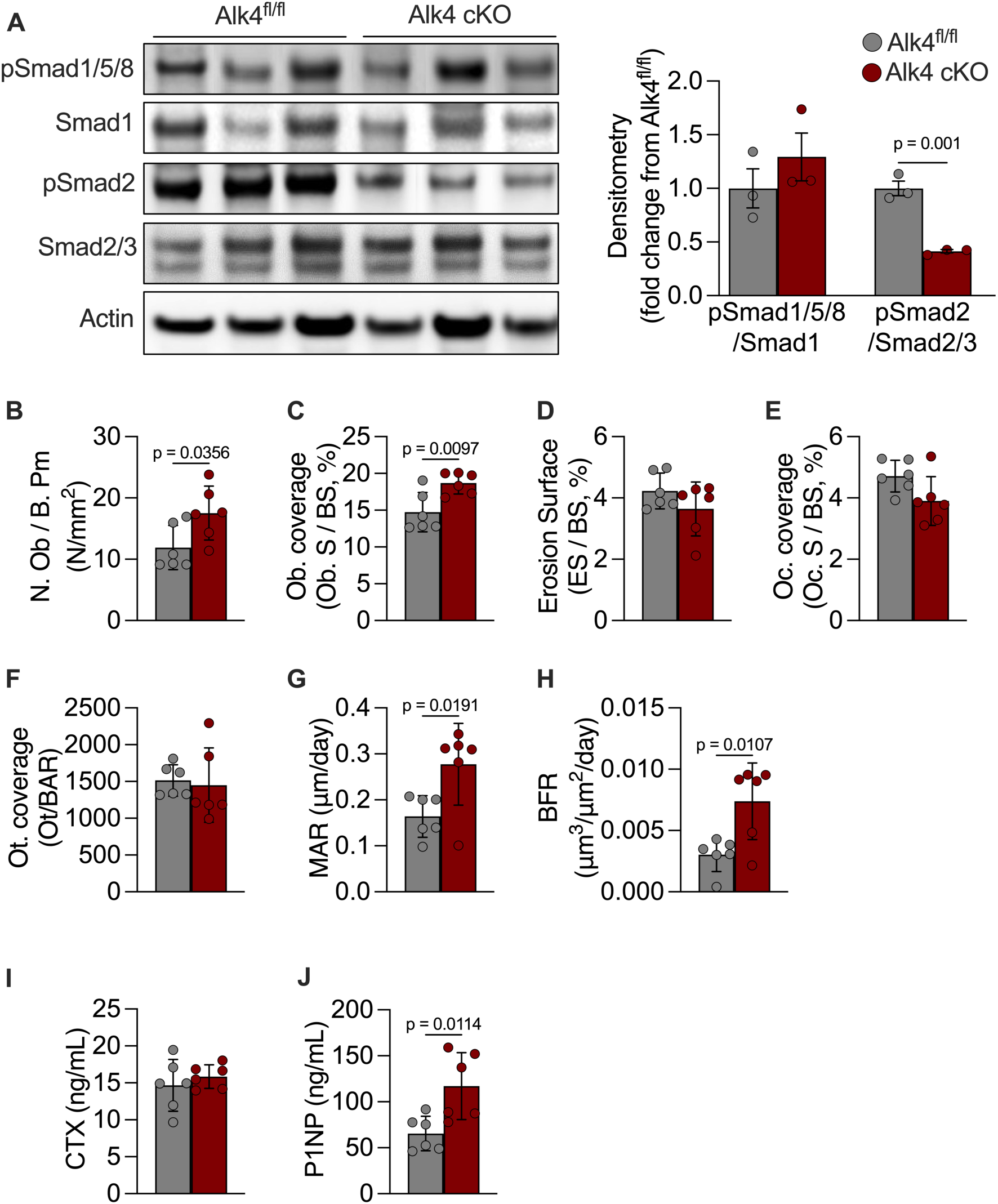
Adult Alk4 cKO mice have reduced levels of pSmad2 and higher osteoblastic activity. Western immunoblots and densitometry analysis (**A**) performed on protein lysates extracted from the marrow compartment of 3-month-old Alk4^fl/fl^ and Alk4 cKO female mice. Histomorphometric measurements were performed in the proximal tibiae of 3-month-old Alk4^fl/fl^ and Alk4 cKO female mice to measure osteoblast number (**B**), osteoblast coverage (**C**), erosion surface (**D**), osteoclast coverage (**E**), Osteocyte coverage (**F**), mineral apposition rate (**G**), and bone formation rate (**H**). Abbreviations: Osteoblasts (Ob.), Bone perimeter (B. Pm), Bone surface (BS), osteoclast (Oc.), osteocyte (Ot.), Mineral apposition rate (MAR), Bone formation rate (BFR). Serum levels of CTX (**I**) and P1NP (**J**) measured by ELISA in 3-month-old Alk4^fl/fl^ and Alk4 cKO female mice. Statistical analyses include unpaired Student *t*-test, p-values shown when < 0.05.

### Deleting Alk4 in primary osteoblasts increases osteogenesis *in vitro*

In Alk4 cKO mice, *Alk4* is deleted in skeletal stem cells and their progeny (osteoblasts and osteocytes), making it difficult to determine if the high bone mass phenotype results from alterations in early progenitors or in differentiated osteoblasts. To address this, we isolated primary osteoblasts from the calvaria of Alk4^fl/fl^ neonates and transfected the cells with adenoviruses expressing either CRE (AdCRE) to delete Alk4 or GFP (AdGFP) to control for the infection. Cells were then cultured in osteogenic conditions for up to 14 days to induce mineralization. *Alk4* RNA expression was successfully decreased in cultures infected with AdCRE (**Supplemental Fig. 2A**). Interestingly, *Alk4* RNA expression gradually increased from 0 to 14 days in osteogenic media as the osteoblasts matured and deposited matrix (**Supplemental Fig. 2A**). After 14 days, we observed increased mineralization marked by elevated alizarin red staining (**Supplemental Fig. 2C**) and alkaline phosphatase staining (**Supplemental Fig. 2C**) in cultures of osteoblasts lacking Alk4 compared to AdGFP controls. Next, we measured whether levels of pSmad1/5/8 or pSmad2 were changed with deletion of *Alk4 in vitro*. Similarly to what was observed in the bones of Alk4 cKO mice, osteoblasts lacking Alk4 had decreased pSmad2 but no change in pSmad1/5/8 levels (**Supplemental Fig. 2D**). Together, our results indicate that *Alk4* RNA expression is upregulated in mature osteoblasts and deletion of *Alk4* increases osteogenesis *in vitro*.

### A subset of osteoblastic genes is regulated by Smad1/5/8 and Smad2/3 in opposite direction

There is ample evidence that bone formation by osteoblasts is enhanced by Smad1/5/8 activation and inhibited by Smad2/3^18, 20, 28–35^. This concept is corroborated by our finding here that reducing levels of pSmad2 without affecting levels of pSmad1/5/8 is sufficient to increase bone formation *in vivo* and *in vitro*. Taken together, these observations prompted us to ask whether the Smad1/5/8 and Smad2/3 pathways compete to regulate the same osteogenic genes. To test this, we performed bulk RNAseq on BMSCs treated with BMP2 and Activin A to activate the Smad1/5/8 and Smad2/3 pathway, respectively. We used W20s, a cell line modelling adult BMSCs that expresses receptors for both BMP2 and Activin A. Cells were cultured with BMP2 or Activin A or a combination of BMP2 + Activin A. After 1hr of treatment, western immunoblots and densitometry analysis confirmed that BMP2 and Activin A treatments induced phosphorylation of Smad1/5/8 and Smad2/3, respectively (**Fig. 5A**). Co-treatment with both ligands increased pSmad1/5/8 and pSmad2 levels but to a lesser degree than individual ligand treatment (**Fig. 5A**).

**Figure 5.**
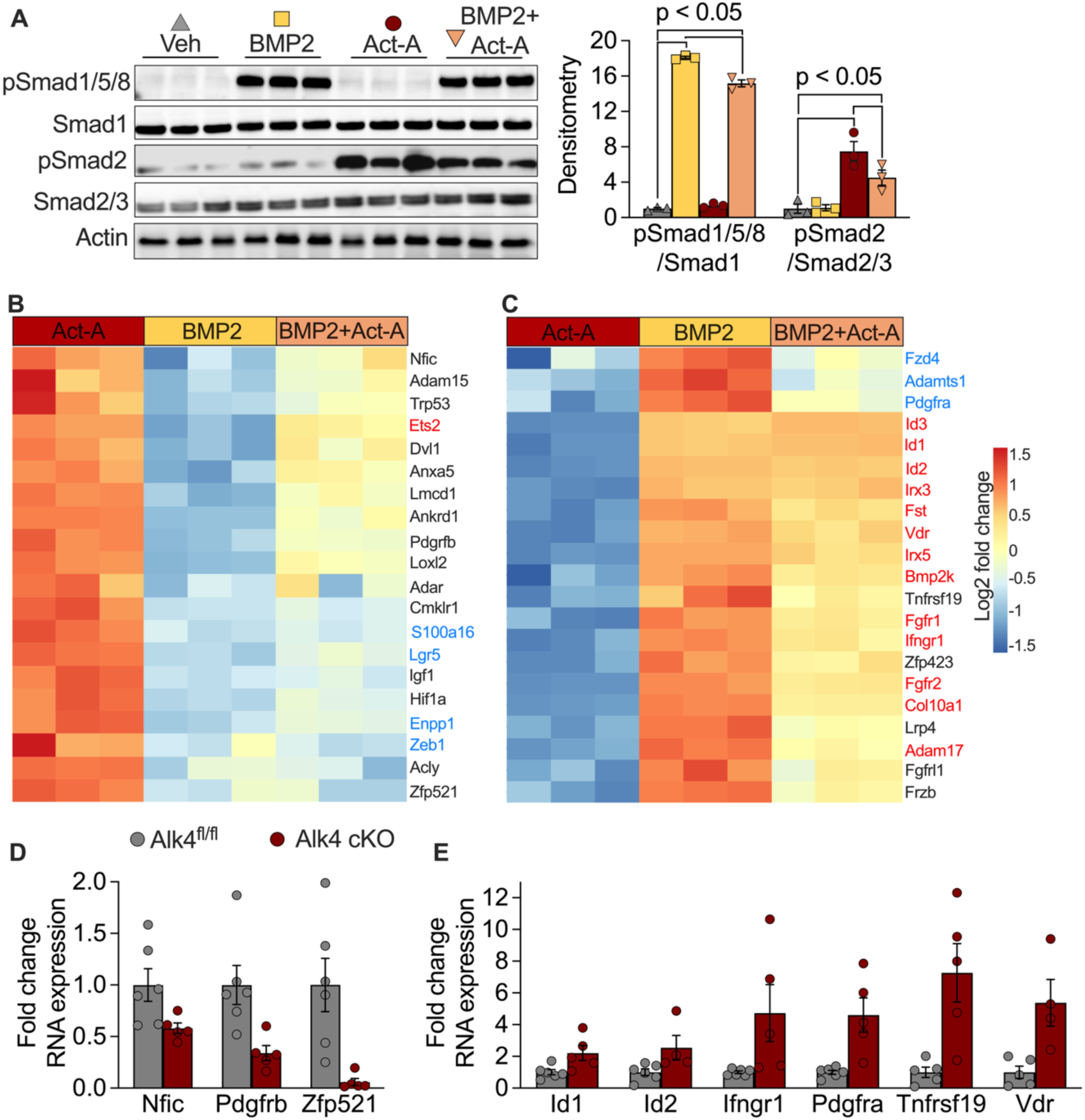
Osteoblastic genes regulated in opposite direction by Smad1/5/8 and Smad2/3. (**A**) Western immunoblots and densitometry analysis (presented as fold change from vehicle) of W20 cultures treated with vehicle PBS (Veh), BMP2 (100ng/mL), Activin A (Act-A, 100ng/mL), or BMP2+Act-A (100ng/mL each). (**B**,**C**) Heatmaps of differentially expressed osteoblastic genes regulated in opposite direction by Activin A and BMP2. Genes in black, blue, and red are either not changed, downregulated, or upregulated by BMP2+Act-A treatment, respectively. qPCR was performed on RNA samples extracted from the femurs of 3-month-old female Alk4^fl/fl^ and Alk4 cKO mice. Genes regulated in opposite direction by Act-A and BMP2 that are significantly (p-value < 0.05) downregulated (**D**) or upregulated (**E**) in the femurs of 2-month-old Alk4^fl/fl^ and Alk4 cKO female mice. Statistical analyses include 2-way ANOVA with multiple comparisons (A) and unpaired Student *t*-test. (D, E).

RNA was extracted for sequencing after 6 hr. of treatment and submitted for sequencing (Raw files and Gene list available at NCBI Gene Expression Omnibus). Using an adjusted p-value cut-off set at 0.05, we identified the differentially expressed genes (DEGs) in each treatment compared to vehicle. Next, we used online gene annotation tools to extract DEGs relevant to osteoblast function and bone formation. Then, we focused on genes regulated in opposing directions by Activin A and BMP2 treatments (**Fig. 5B, C**). We found 20 osteoblastic genes upregulated by Activin A and downregulated by BMP2 (**Fig. 5B**) and 21 osteoblastic genes downregulated by Activin A and upregulated by BMP2 (**Fig. 5C**). When cells were co-treated with BMP2+Activin A, out of these 41 genes, 20 had no detectable change in their expression, 17 followed the pattern of expression of BMP2 treatment, and 4 followed the pattern of expression of Activin A treatment (**Fig. 5B, C**). Together, our data demonstrate that 41 genes relevant to osteoblast function and bone formation are regulated in opposite direction by Smad1/5/8 and Smad2/3. Our results also demonstrate that osteogenic cells respond to co-treatment with Activin A and BMP2 by producing a combinatorial transcriptional response.

Finally, we considered whether some of these 41 genes adopt a Smad1/5/8-like pattern of expression in the femurs of Alk4 cKO mice (where Smad2/3 activity is decreased). We extracted RNA from femurs of 3-month-old Alk4^fl/fl^ and Alk4 cKO female mice and performed qPCR. We found significantly decreased expression of *Nfic*, *Pdgfrb*, and *Zfp521* in Alk4 cKO mice compared to Alk4^fl/fl^ controls (**Fig. 5D**). *Nfic* and *Zfp521* encode transcription factors involved in regulating osteogenic cell differentiation^36–42^. PDGFRβ, the receptor for PDGF-BB, is expressed on the surface of osteoblasts and its effects on bone formation are linked to the presence of PDGFRα^43, 44^. *Pdgfra*, along with *Id1*, *Id2*, *Ifngr1*, *Tnfrsf19*, and *Vdr* had increased expression in the bones of Alk4 cKO mice compared to Alk4^fl/fl^ controls (**Fig. 5E**). *Id1* and *Id2* are direct target genes of Smad1/5/8 and are known to be upregulated during BMP-induced osteoblast differentiation^45–49^. *Ifngr1* encodes for the receptor to IFNγ, a cytokine that promotes mineralization by osteoblasts in a time-specific manner and *Ifngr1* null mice exhibit low bone volume and bone turnover akin to osteoporosis^50–52^. *Tnfrsf19* also encodes for a receptor, and its locus is associated with BMD in humans^53^. Additionally, *Tnfrsf19* is upregulated by Wnt3a treatment to block the differentiation of human mesenchymal stem cells into adipocytes and promote alkaline phosphatase activity, consistent with a pro-osteogenic effect^54^. Finally, the vitamin D receptor (*Vdr*), also a transcription factor, is crucial for the differentiation and mineralization function of osteoblasts^55–58^. Collectively, these findings suggest that when Alk4 is deleted in skeletal stem cells, a subset of genes central to osteoblast function and differentiation adopts a pattern of expression akin to that induced by the Smad1/5/8 pathway.

### Treatment with Alk4-Fc increases bone volume and muscle mass in older mice

We next asked whether ligands that bind Alk4 could be therapeutically targeted to alleviate age-related bone loss in older mice. To test this, we designed a preclinical study using a drug consisting of the fragment crystallizable (Fc) of Alk4 (Alk4-Fc). Theoretically, Alk4-Fc binds circulating Alk4-binding ligands and neutralizes them from the circulation. A similar Alk4-Fc drug was previously shown to increase BMD and muscle mass in young mice by capturing circulating activins^59, 60^. We injected 24-month-old C57BL6J male mice with Alk4-Fc for 2 months and measured BMD and lean mass by DXA weekly. Alk4-Fc treatment did not significantly affect body weight (**Fig. 6A**) or fat mass (**Fig. 6B**), over the course of the study. Although lean mass was also not changed significantly by Alk4-Fc treatment, we noted a mild increase in Alk4-Fc-treated mice after 5 weeks of treatment (**Fig. 6C**). BMD was significantly increased in Alk4-Fc -treated mice compared to controls by the end of the experiment (**Fig. 6D**). We then harvested muscles from the mice and measured their weight at dissection. Gastrocnemius (**Fig. 6E**), quadriceps (**Fig. 6F**), and triceps (**Fig. 6G**) isolated from Alk4-Fc -treated mice had higher weights than those from controls. To investigate effects on bone volume, we performed microCT on femurs and L5 vertebrae. Trabecular bone volume (**Fig. 6H**) and BMD (**Fig. 6I**) were increased in the femurs of mice treated with Alk4-Fc. As observed in Alk4 cKO mice (**Fig. 3G**), we did not detect any change in cortical area with Alk4-Fc injections (**Fig. 6J**). Vertebral trabecular bone volume (**Fig. 6K**) and BMD (**Fig. 6L**) were also significantly higher in Alk4-Fc treated mice compared with controls. Finally, we measured circulating Activin A levels at the end of the study and saw a significant reduction in Alk4-Fc treated animals (**Fig. 6M**). Our results demonstrate that Alk4-Fc treatment alleviate age-related muscle and bone loss by reducing levels of circulating Activin A in old mice.

**Figure 6.**
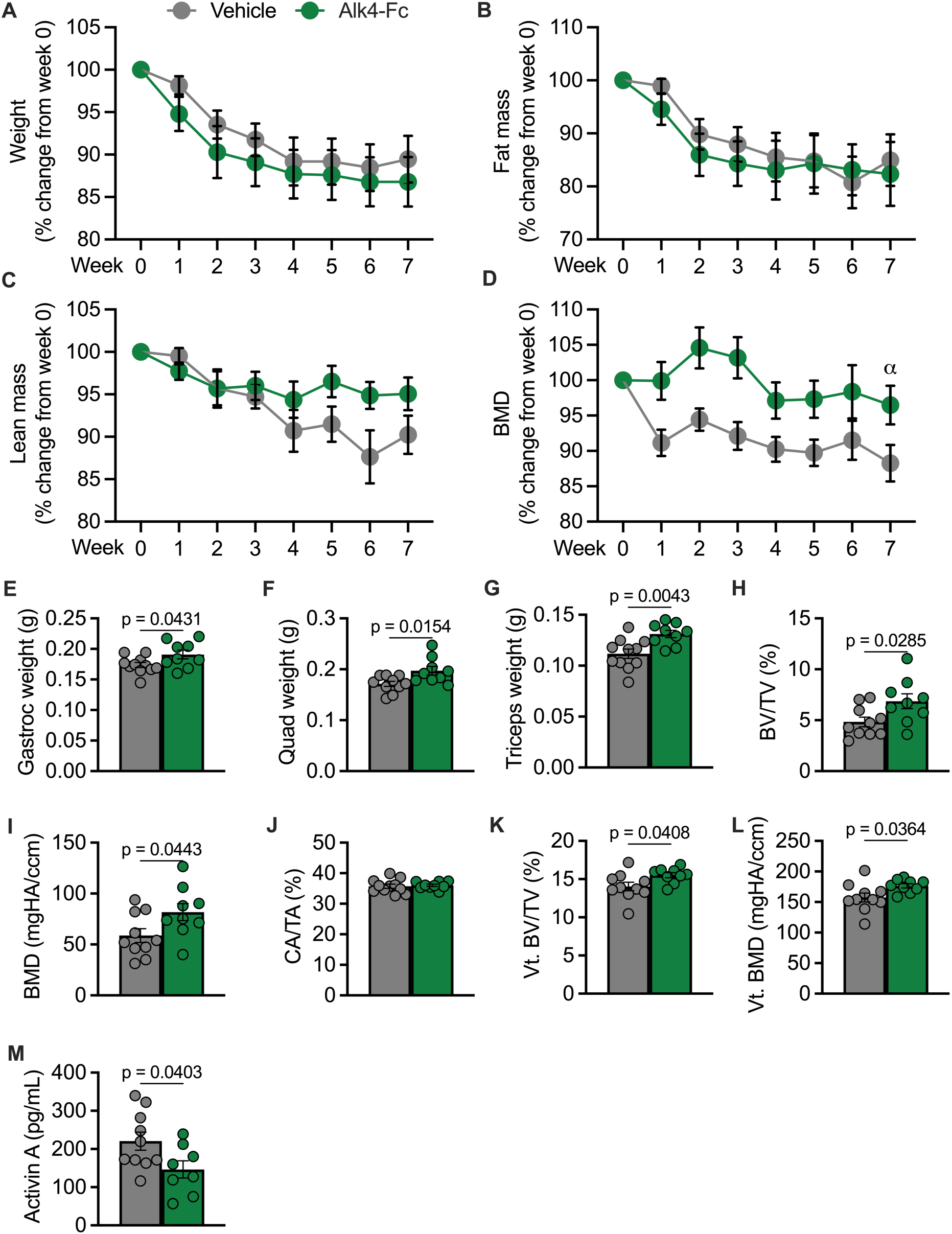
Alk4-Fc treatment protects against bone and muscle loss in old mice. 24-month-old C57BL6J male mice were injected with 5mg/kg body weight of Alk4-Fc or vehicle (PBS). Weekly DXA measurements (as % change from baseline) for total body weight (**A**), fat mass (**B**), lean mass (**C**), and BMD (**D**) over the course of the study. α denotes p-value < 0.05 when running Student *t*-test on BMD endpoint measurements. Endpoint muscle weights for gastrocnemius (gastroc, **E**), quadriceps (quad, **F**), and triceps (**G**). Endpoint microCT analyses were performed in distal femurs to measure trabecular bone volume (bone volume /total volume: BV/TV, **H**) and trabecular BMD (**I**). Cortical area fraction (cortical area / total area: CA/TA, **J**) was measured at the midshaft diaphysis. For vertebral (Vt.) bone, analysis was performed in the L5 vertebral body to measure trabecular bone volume (**K**) and trabecular BMD (**L**). Circulating levels of Activin A (**M**) were measured by ELISA in sera samples at the end of the study. Statistical analyses include unpaired Student *t*-test, p-values shown when < 0.05.

## DISCUSSION

pending

## METHODS

### Animals

Mice were housed in cages with sterilized paper bedding in a 14:10-hour light/dark cycles with ad libitum access to sterilized water and chow. Alk4 floxed mice (Alk4^fl/fl^) on the C57BL/6J background were donated by Dr. Gloria Su who previously described the generation of that allele^76^. Alk4^fl/fl^ mice were bred to Prx1-Cre (B6.Cg-Tg(Prrx1-cre)1Cjt/J) obtained from The Jackson Laboratory (Strain no. 005584) to produce Alk4 cKO mice. All C57BL6J mice were obtained from The Jackson Laboratory. Figures contain data combining both male and female mice before 15 days of age. After 15 days of age, female and male data are presented separately in figures.

### Study approval

All experiments involving animals were performed in compliance with the Guide for the Care and Use of Laboratory Animals and were approved by the Harvard Medical Area Institutional Animal Care and Use Committee.

### Western Immunoblots

Protein samples were collected from bone marrow flushed from bones or from cell cultures. Proteins were extracted using RIPA lysis buffer (Cell Signaling Technology) supplemented with Haly Protease & Phosphatase Inhibitor Cocktail (Fisher Scientific). Protein lysates were resolved on 4-12% gradient Bis-tris Plus gels (Fisher Scientific) and transferred on nitrocellulose membranes. Membranes were blocked using 5% non-fat dry milk, 5% bovine albumin serum in TBST. Blotting was performed using the following primary antibodies: phospho-Smad1/5/8 (Cell Signaling Technology, #13820), phospho-Smad2 (Cell Signaling Technology, #3108), Smad1 (Cell Signaling Technology, #9743), Smad2/3 (Cell Signaling Technology, #8685), and βactin (Cell Signaling Technology, #4967). Secondary blotting was performed using anti-rabbit IgG conjugated to horseradish peroxidase (HRP) antibody (Cell Signaling Technology, #7074). Blots were developed using SuperSignal West Femto Substrate (Fisher Scientific). Signal intensities were quantified by ImageJ software to produce densitometry plots. Each band signal was normalized to its actin band to account for loading variation. The ratios of pSmad1/5/8/Smad1 and pSmad2/Smad2/3 were calculated and reported on densitometry plots.

### Quantitative PCR (qPCR)

RNA samples were collected from femurs and cell cultures and converted to cDNA as previously described ^77^. qPCR was performed using the following mouse primers: *Acvr2a* (F: CCCTCCTGTACTTGTTCCTACTCA, R: GCAATGGCTTCAACCCTAGT); *Acvr2b* (F: ATCGTCATCGGAAGCCTCCC, R: CAGCCAGTGATCCTTAATC); *Actb*: (F: GTGTACGACCAGAGGCATAC, R: AAGGCCAACCGTGAAAAGAT); *Alk1 (F:* CGGCTCTGGACGTGAGAC, R: GCAAAACGTGATAGCTGTGG); *Alk2 (F:* ACCCTGTTGGAGTGTGTCG, R: CGTCTCCCTGAACCATGACT); *Alk3 (F:* AACAGCGATGAATGTCTTCGAG, R: GTCTGGAGGCTGGATTATGGG); *Alk4* (F: TTCTTCCCCCTTGTTGTCCTC, R: ACAGGTGTAGTTGGTCTGTAGG); *Alk5* (F: CAGCTCCTCATCGTGTTGG, R: CAGAGGTGGCAGAAACACTG); *Alk6 (F:* CCCTCGGCCCAAGATCCTA, R: CAACAGGCATTCCAGAGTCATC); *Alk7 (F:* ATGCAGGAAAAACACTCAAGG, R; GAACCAAGAGAGGCAGACCA); *Bmp2 (F:* AGATCTGTACCGCAGGCACT, R: GTTCCTCCACGGCTTCTTC); *Bmp4 (F:* GAGGAGTTTCCATCACGAAGA, R: GCTCTGCCGAGGAGATCA); *Bmp6 (F:* ACTGACTAGCGCGCAGGA, R: TGTGGGGAGAACTCCTTGTC); *Bmp7 (F:* CGAGACCTTCCAGATCACAGT, R: CAGCAAGAAGAGGTCCGACT); *Bmpr2* (F: GAGCCCTCCCTTGACCTG, R: GTATCGACCCCGTCCAATC); *Id1 (F:* GCGAGATCAGTGCCTTGG, R: CTCCTGAAGGGCTGGAGTC); *Id2 (F:* GACAGAACCAGGCGTCCA, R: AGCTCAGAAGGGAATTCAGATG); *Ifngr1 (F:* CTTGAACCCTGTCGTATGCTGG, R: TTGGTGCAGGAATCAGTCCAGG); *Inhba (F:* ATCATCACCTTTGCCGAGTC, R: TCACTGCCTTCCTTGGAAAT); *Inhbb (F:* GATCATCAGCTTTGCAGAGACA*, R:* TGCCTTCATTAGAGACGAAGAA); *Nfic (F:* GACCTGTACCTGGCCTACTTTG, R: CACACCTGACGTGACAAAGCTC); *Pdgfra (F:* TGCAAATTGACATAGAAGGAGAAG, R: GCCCTGTGAGGAGACAGC); *Pdgfrb (F:* TCAAGCTGCAGGTCAATGTC, R: CCATTGGCAGGGTGACTC); *Tgfb1 (F:* TGGAGCAACATGTGGAACTC, R: GTCAGCAGCCGGTTACCA); *Tgfbr2* (F: GGCTCTGGTACTCTGGGAAA, R: AATGGGGGCTCGTAATCCT); *Tnfrsf19 (F:* TGTGTCCTCTGCAAACAGTGCG, R: CCAGTCTTCCTTGAACCGGTGC); *Vdr (F:* GCTTCCACTTCAACGCTATGA, R: ATGCGGCAATCTCCATTGAA); *Zfp521 (F:* ACAACGAGTGGGACATCCAGGT, R: GCTGTGCTCTATCAGGTGACAC). Data were normalized using *Actin* (*Actb*) as a housekeeping gene and analyzed using the ΔΔCT method.

### Serum ELISA

Quantitative determination of serum N-terminal propeptide of type I procollagen (P1NP) and C-terminal telopeptide of type 1 collagen (CTX-1) were performed using commercial kits (Mouse P1NP EIA and RatLaps CTX-1 EIA, Immunodiagnostic Systems). Serum Activin A and TGFβ1 levels were determined by Quantikine ELISA Kits (R&D Systems, Activin A #DAC00B, TGFβ1 #DB100C). Assays were performed according to manufacturer’s instructions.

### Skeletal preparation and histology analysis of newborn mice

Newborn mice were prepared and stained with Alizarin Red and Alcian Blue to identify mineralized bone and cartilage as previously described ^78^. Limbs from newborn mice were dissected and fixed in 10% neutralized buffered formalin, paraffin embedded, and sectioned. Slides were stained with Toluidine Blue.

### Micro-computed tomography (MicroCT) analysis

Femurs and L5 vertebrae were collected from mice and fixed overnight in 10% neutralized buffered formalin. Samples were scanned using a μCT35 (Scanco Medical AG) and analysis was performed according to guidelines published by the American Society for Bone and Mineral Research^79^.

### Static and dynamic histomorphometric analyses

For dynamic measurements, mice were injected intra-peritoneally with calcein (20mg/kg of body weight) 7 and 2 days prior to sacrifice. Tibias were collected and fixed in 10% neutralized buffered formalin. Processing and bone histomorphometric analyses were performed on the proximal tibia by the Histology & Histomorphometry Services offered by the Center for Skeletal Research (Boston, MA) according to guidelines published by the American Society for Bone and Mineral Research ^80^.

### Cell culture

Primary bone marrow stromal cells from the femurs and tibias of 8-week-old C57BL/6J female mice were isolated and cultured as previously described ^81^. Cultures were treated with sera in presence or absence of 1μg/mL of anti-TGFβ1,2,3 antibody (Bio-techne, #MAB1835). Primary osteoblasts were isolated from the calvariae of newborn Alk4^fl/fl^ mice as previously described ^82^. Osteoblasts were cultured in basal medium (DMEM containing 10% FBS and supplemented with 100 U/ml penicillin-G and 100 mg/ml streptomycin, Gibco). Knockdown of Alk4 in osteoblasts was achieved by Adenovirus infection using Ad5-CMV-eGFP (AdGFP) and Ad5-CMV-Cre-eGFP (AdCRE) viral particles obtained from the University of Iowa Gene Transfer Vector Core. Osteoblasts were seeded at 400,000 cells/well in 24-well plates overnight and transduced with adenoviruses for 6 hours. For osteoblast differentiation, confluent cultures were placed in osteogenic media constituting of basal medium supplemented with 0.2 mM ascorbic acid and 10 mM β-glycerophosphate. Alkaline phosphatase staining was performed using the Leukocyte Alkaline Phosphatase kit (Sigma-Aldrich) according to the manufacturer’s guidelines. Alizarin red staining was performed by incubating fixed cells with 1% Alizarin Red S, pH 4.1-4.3 for 20min at room temperature. Area of staining was quantified using ImageJ software. W20 cells were cultured in basal medium until confluent. Cultures were deprived of serum for 12 hours before being treated with vehicle (PBS), 100ng/mL of BMP2, 100ng/mL Activin A (R&D Systems, #338-AC-010/CF), or a combination of both BMP2 + Activin A (100ng/mL each). After 1hr of, proteins were extracted for western immunoblotting. After 6hr, RNA samples were collected for RNA Sequencing.

### RNA Sequencing

RNA from each sample was sent to the Biopolymers Facility at Harvard Medical School to determine RNA quality. Samples that met or exceeded the RiN value threshold of 7 (Agilent 2100 BioAnalyzer) were were sent to BGI Genomics (Beijing, China) to construct libraries and perform sequencing. For library construction, poly-A+ mRNA molecules were isolated, and cleaved using divalent cations under elevated temperature. Next, cleaved RNA fragments were copied into first strand cDNA using reverse transcriptase and random primers. This was followed by second strand cDNA synthesis using DNA Polymerase I and RNase H. The cDNA fragments then had the addition of a single ‘A’ base and subsequent ligation of the adapter. The resulting products were purified and enriched with PCR amplification. The PCR yield was then quantified by Qubit and samples were pooled together to make a single strand DNA circle (ssDNA circle), which gave the final library. DNA nanoballs (DNBs) were generated with the ssDNA circle by rolling circle replication to enlarge the fluorescent signals at the sequencing process. The DNBs were loaded into the patterned nanoarrays and pair-end reads of 100 bp were read through on the BGISEQ-500 platform for the following data analysis study. For this step, the BGISEQ-500 platform combines the DNA nanoball-based nanoarrays and stepwise sequencing using Combinational Probe-Anchor Synthesis Sequencing Method. After filtering, the resulting reads were stored in FASTQ format. FASTQ files were sent to the Harvard Chan Bioinformatics Core for analysis. All samples were processed using a bulk RNA-seq pipeline implemented in the bcbio-nextgen project (bcbio-nextgen,1.2.8-1c563f1). Gene expression was quantified using Salmon (version 0.14.2) ^83^. To determine which genes were differentially expressed between the vehicle and the three treatments, DESeq2 (version 1.30.1) ^84^. An adjusted p-value cut-off of 0.05 was applied to identify differentially expressed genes.

### Alk4-Fc treatment

Alk4-Fc was obtained from Bio-Techne (Minneapolis, MN). The drug was manufactured in a single sterile batch, lyophilized, and stored in 1mg aliquots at -80°C. Resuspension in PBS was made on the days of injection. 23-month-old C57BL6/J male mice were ordered from The Jackson Laboratories. An acclimatation period of 1 month elapsed before the study started. A volume up to 150μL was injected twice a week intraperitoneally to yield a dose of 5mg/kg of body weight (n=12 mice). The same volume of PBS was used for vehicle controls (n=12 mice). Body weight was measured weekly to adjust the dose of Alk4-Fc. The study lasted 8 weeks before mice were euthanized for tissues and fluid collection. By the end of the study, 5 mice were found dead (n=2 in the control group, n=3 in the Alk4-Fc treated group). A full body necropsy was performed on each animal found dead and cause of death was determined to be natural.

### Dual X-ray Absorptiometry

Mice were scanned once a week using the iNSiGHT system which employs a combination of a low energy (60 kV, 0.80 mA) and a high energy (80kv, 0.80 mA) scan (Osteosys, Seoul, Korea). Before each scanning session, the machine was calibrated using a standard phantom block. Each mouse was anesthetized using isoflurane before being positioned and scanned. Measurements were calculated by the iNSiGHT software excluding the head of the animal to prevent effects from the teeth and ear tags. Measurements included BMD (g/cm2), lean mass (g), fat mass (g), fat proportion (%). For each measurement, data collected at the beginning of the study (Week 0) was used as a baseline to calculate % change for each mouse over the duration of the study.

### Statistics

The data are expressed as the mean ± standard error of the mean (SEM). Results were analyzed for statistically significant differences using Prism 8 statistical software (GraphPad Software, Inc., La Jolla, CA, USA). We performed Student’s t test or ANOVA with multiple comparisons with statistical significance set at p < 0.05. For mice cohorts, we powered the study (β = 0.95) to complete all assays with a standard deviation of 10% for an α (p < 0.01). Numbers of mice (n) used in specific experiments are indicated in each figure.

## Author contributions

DEM, SMC, LG, and VR conceived the project and designed the research studies. Data collection was performed by DEM, SMC, KMB, EVD, KK, EK, and KA. Analysis and interpretation of the data was conducted by DEM, LG, VR and RSK. DEM, LG and VR composed the manuscript. VR managed the funding source for the research project. All the authors read, edited and approved the final version of the manuscript.

## Acknowledgements

The authors thank Dr. Gloria Su for donating the Alk4^fl^ mice. We are grateful for the Histology & Histomorphometry Services offered by the Center of Skeletal Research in Boston, MA. The authors would like to thank Radhika S. Khetani of the Harvard Chan Bioinformatics Core, Harvard T.H. Chan School of Public Health, Boston, MA for performing the bulk RNA-seq analysis. This work was partially funded by the Harvard Medical School Foundry. This work was supported by the NIH grant 5R01AR064227.

---

**Supplemental Figure 1.**
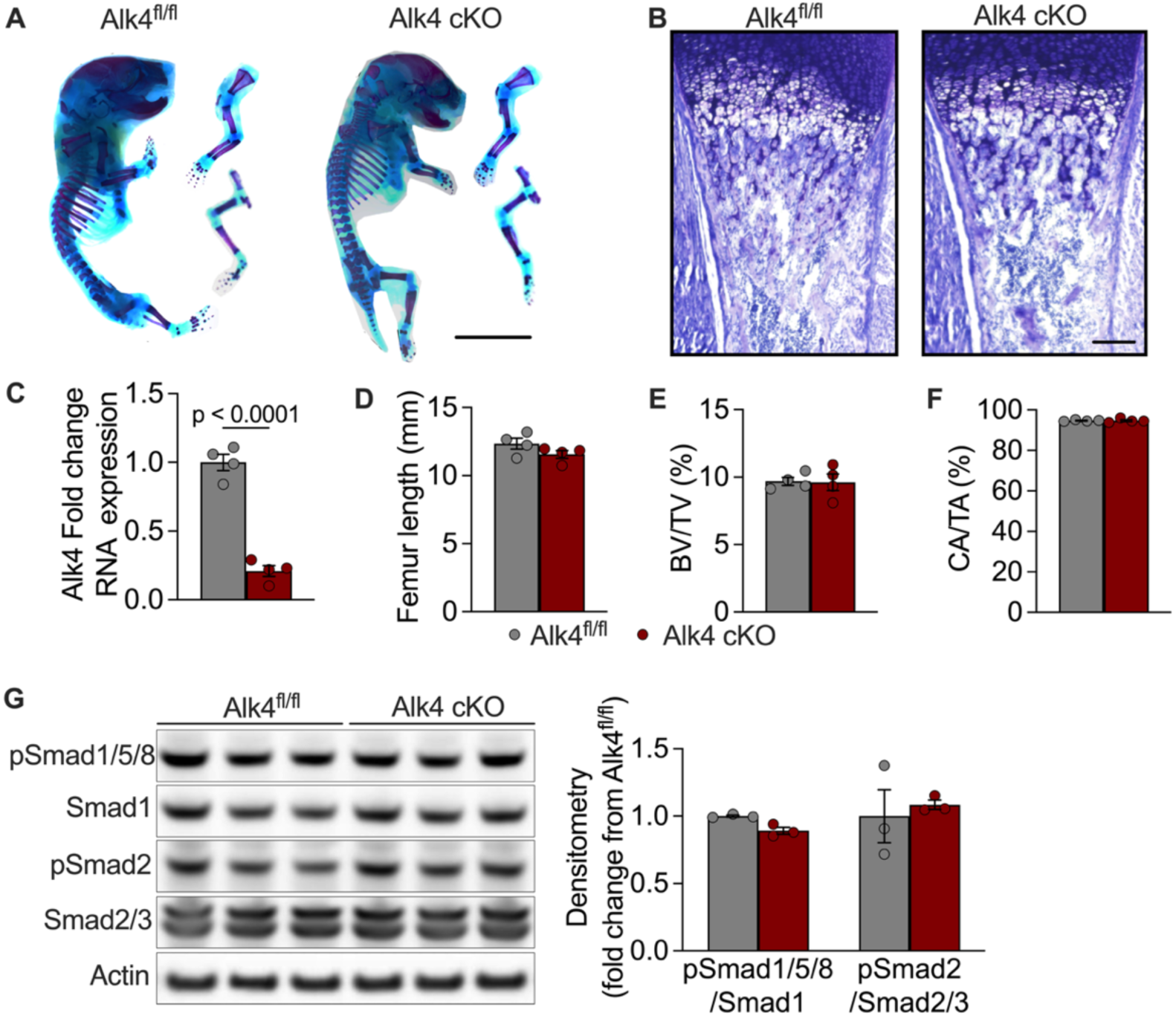
Skeletogenesis and early postnatal bone formation are not affected in young Alk4 cKO mice. (**A**) Whole mount skeletal preps of hindlimbs and (**B**) toluidine blue sections of distal femurs of postnatal day 0 (P0) Alk4^fl/fl^ and Alk4 cKO mice. Scale = 0.5mm. (**C**) qPCR measurement of *Alk4* RNA expression in femurs of 4-week-old Alk4^fl/fl^ and Alk4 cKO mice. (**D**) Femur length, (**E**) bone volume / total volume (BV/TV), and (**F**) cortical area / total area (CA/TA) obtained by microCT performed on femurs of 4-week-old Alk4^fl/fl^ and Alk4 cKO mice. (**G**) Western immunoblotting and densitometry analysis performed on protein lysates extracted from marrow-free femurs of 4-week-old Alk4^fl/fl^ and Alk4 cKO mice. Statistical analyses include unpaired Student *t*-test, p-values shown when < 0.05.

**Supplemental Figure 2.**
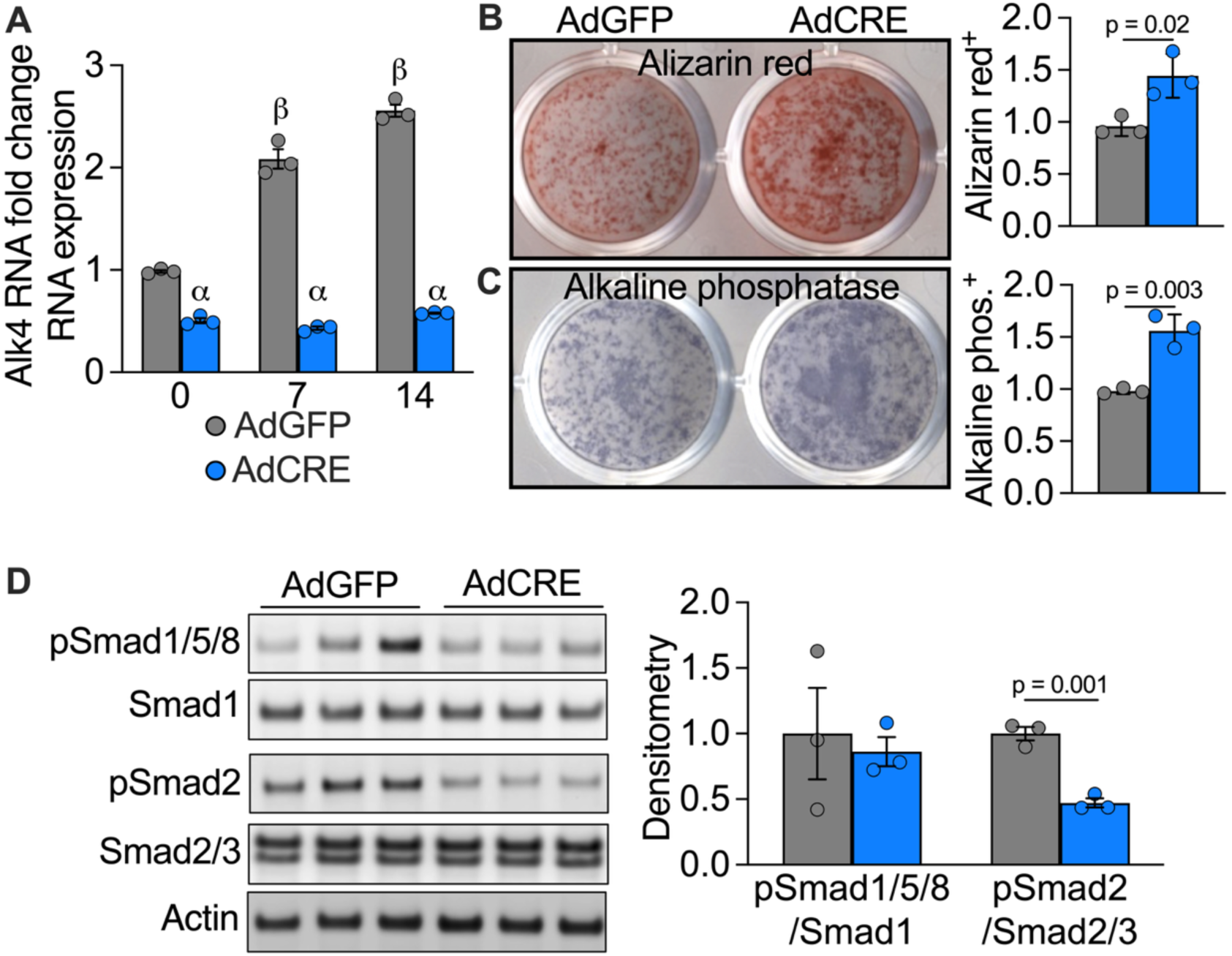
Reduced pSmad2 levels and increased osteogenesis in osteoblasts lacking Alk4 *in vitro*. Primary calvarial osteoblasts isolated from neonates Alk4^fl/fl^ mice were transfected with adenoviruses containing GFP (AdGFP) or CRE (AdCRE). (**A**) qPCR measurement of *Alk4* RNA expression in primary calvarial osteoblasts cultured in osteogenic media for 14 days. (**B**) Alizarin red and (**C**) alkaline phosphatase staining of primary calvarial osteoblasts after 14 days in osteogenic media. Quantification of stained area is presented as fold change from AdGFP. (**D**) Western immunoblots and densitometry analysis performed on protein lysates extracted from cultures after 14 days in osteogenic media. Densitometry analysis is presented as fold change from AdGFP. Statistical analyses include: 2-way ANOVA with multiple comparisons (A, D) with α showing adjusted p-value <0.05 when comparing between genotypes at the same time point and β showing adjusted p-value <0.05 when comparing from a time point to the previous one within the same genotype; and Student *t*-test (B, C), p-values shown when < 0.05.

## Notes

### Competing Interest Statement

The authors have declared no competing interest.

